# Structural determinants of relaxation dynamics in chemical reaction networks

**DOI:** 10.1101/2022.05.25.493374

**Authors:** Yusuke Himeoka, Julius B. Kirkegaard, Namiko Mitarai, Sandeep Krishna

## Abstract

Understanding the relationship between the structure of chemical reaction networks and their reaction dynamics is essential for unveiling the design principles of living organisms. However, while some network-structural features are known to relate to the steady-state characteristics of chemical reaction networks, mathematical frameworks describing the links between out-of-steady-state dynamics and network structure are still underdeveloped. Here, we characterize the out-of-steady-state behavior of a class of artificial chemical reaction networks consisting of the ligation and splitting reactions of polymers. Within this class, we examine minimal networks that can convert a given set of inputs (e.g., nutrients) to a specified set of targets (e.g., biomass precursors). We find three distinct types of relaxation dynamics after perturbation from a steady-state: exponential-, power-law-, and plateau-dominated. We computationally show that we can predict this out-of-steady-state dynamical behavior from just three features computed from the network’s stoichiometric matrix, namely, (i) the rank gap, determining the existence of a steady-state; (ii) the left null-space, being related to conserved quantities in the dynamics; and (iii) the stoichiometric cone, dictating the range of achievable chemical concentrations. We further demonstrate that these three quantities also predict the type of relaxation dynamics of combinations of our minimal networks, larger networks with many redundant pathways, and a real example of a metabolic network. The unified method to predict the qualitative features of the relaxation dynamics presented here can provide a basis for understanding the design of metabolic reaction networks as well as industrially useful chemical production pathways.

**Author summary:** The relationship between network structure and chemical reaction dynamics is of central interest in chemical reaction network theory, as it underlies chemical manufacturing, cellular metabolism, and bioengineering. The links between structure and steady-state properties have been extensively investigated. However, how far the network structure determines the out-of-steady-state, transient dynamics of chemical reactions is unexplored. Here we construct a chemical reaction network model that is simple but generates a wide variety of network instances. By computationally exploring the networks’ structural- and dynamical features, we found that three network-structural features are sufficient to predict the qualitative characteristics of the relaxation dynamics after the chemical concentrations are perturbed from their steady-state. Depending on the values of those three features, the chemical reaction dynamics on the network exhibit exponential, plateau, and power-law relaxation. Also, we found that such features are determinants of the dynamics of biological metabolic reaction systems. Our findings provide a foundation for the structure-based prediction of chemical reaction dynamics.

## Introduction

Origins of chemical reaction network theory can be found in the discovery of the law of mass action [1]. Since then, mathematical laws governing chemical reactions have been uncovered, such as Le Chatelier’s principle [2], the Arrhenius equation [3], and the detailed-balance condition for the feasibility of chemical equilibrium [4].

For the development of quantitative theories of living entities, a deep understanding of the out-of-steady-state dynamics of chemical reactions is indispensable. Until recently, most experimental studies of microbial physiology focused on steadily-growing cellular populations. In such situations, the assumption that the concentrations of cellular metabolites are constant (at steady-state) is reasonable, and static metabolic analysis such as Flux Balance Analysis [5–10] and steady-state cell models [11, 12] have provided a useful mathematical description of stable cell growth.

However, recent experiments have been unveiling the dynamic nature of cellular physiology more quantitatively. For instance, in starved *Escherichia coli*, the lag time — the duration that the cell takes to restart growth after the substrate replenishment — was shown to depend on how long the cells are starved [13–17] and the death rates of the starved cells differ depending on the previous culture conditions, even though the starvation condition is identical [18]. Such “memory” is a unique feature of out-of-steady-state systems. In addition, the time scales of the memories are much longer than that in steadily-growing cells. It is therefore desirable to develop theoretical frameworks to understand how such slow relaxation dynamics and long-term memory emerge from the structure of chemical networks.

There are two broad types of approaches one might take to explore the link between the network structure and the relaxation characteristics of chemical networks. One is to examine random networks with structural characteristics (such as density of links and degree distribution) identical to natural chemical networks like metabolic networks [19–22]. This approach enables us to explore a large ensemble of network structures, but due to the simplification into a random network, it may miss crucial properties of real chemical networks. For example, the central principle of chemical reactions, the law of mass conservation, may be violated. Thus, any structure-dynamics connections uncovered by these approaches may not always apply to real chemical networks.

On the other hand, data-driven approaches such as reaction pathway prediction by using machine learning [23–25] generate realistic chemical reaction networks, which never violate mass conservation and other necessary properties. However, it is too computationally costly to explore a wide variety of possible chemical reaction networks. The generality of structure-dynamics connections, or lack of it, uncovered by these approaches may therefore be unclear.

Here we develop an alternative approach. We construct a class of the chemical reaction network consisting of chemical species that can be represented as monomer sequences (polymers), and chemical reactions that ligate (concatenate) and split (fragment) these polymers. The model can generate a wide variety of chemical reaction network structures that are guaranteed to satisfy the mass conservation principle.

Microbial metabolic networks and chemical networks used in industrial manufacturing processes have the specific function of converting certain input chemicals into target chemicals. For instance, in bacterial metabolism, carbon, nitrogen, phosphorous sources (e.g., glucose, potassium hydrogen phosphate, ammonium chloride) and other miscellaneous molecules are converted into biomass such as proteins, nucleic acids, and lipids. Therefore, we impose a further constraint on the networks we study - the ability to synthesize a given set of targets from a specified set of input chemicals. The resultant chemical reaction networks we generate are, of course, a subset of all possible networks, but they form a well-defined subset so the generality of the structure-dynamics relationships we find is also well-defined. Our framework can also easily be extended to include other kinds of chemical species and reactions as desired.

We find that for these artificial reaction networks we construct, there is a single stable steady-state for the chemical concentrations. Computational simulations of the chemical reaction dynamics upon perturbation of these steady-states revealed four distinct types of relaxation dynamics. Interestingly, we found that the qualitative features of the relaxation dynamics are determined by only three properties of the stoichiometric matrix which encodes the chemical network structure, namely, (i) the rank gap, (ii) the dimension of the left null space, and (iii) the stoichiometric cone. We will define all these properties precisely in subsequent sections, but crudely, a non-zero rank gap indicates that no steady-state exists in the absence of any degradation of chemicals, and the left null space determines the existence of conserved quantities in the dynamics. The stoichiometric cone determines the reachability of the final steady state. Finally, we tested the applicability of these three network-structural features to the prediction of relaxation dynamics of a real biological (metabolic) network. Our results and the general framework we have developed provides valuable tools for exploring the relationship between the network structure and chemical reaction dynamics, which may be applied both to complex industrial manufacturing processes as well as design of synthetic microorganisms designed to perform certain chemical tasks [26, 27].

## Model

### Constructing a class of artificial reaction networks

For the construction of the chemical reaction networks, we define a set of chemicals 𝒞 and the set of the reactions ℛ they participate in. In our artificial reaction network model, we consider each chemical species to be a one-dimensional chain of the monomers, such as *AA* or *AABA* where *A* and *B* represent different monomer species. Each monomer idealizes chemical elements such as hydrogen, carbon, nitrogen, and oxygen, or even larger chemical moeities. The chemical reactions possible are the splitting or ligation of the chemicals. For instance, there are three splitting reactions possible with *AABA* as the substrate: *AABA* →*A* + *ABA, AABA* →*AA* + *BA*, and *AABA* →*AAB* + *A*. In the present manuscript, we suppose that all reactions are reversible. That is, if there is a splitting reaction *AABA* → *A* + *ABA*, the corresponding ligation reaction *A* + *ABA* → *AABA* exists.

With this set-up, the set of all possible chemicals 𝒞 and the set of all possible reactions ℛ are fully determined by setting the number of monomers *M* and the length of the longest polymer *L*. For instance, in the case of *M* = 2 and *L* = 2, 𝒞 and ℛ are given by

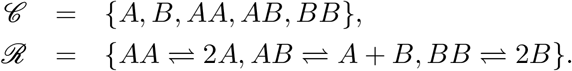

To avoid redundancy, we omit the direction of the polymers, e.g., the polymer *BA* is identified as identical to *AB*. Given this, the number of chemicals (|𝒞 |) and the number of reactions (|ℛ|) scale with the parameters *M* and *L* as

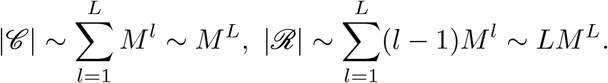

We now construct individual chemical reaction networks by choosing appropriate subsets of ℛ. Importantly, as each reaction in ℛ satisfies the mass-conservation principle, all the reaction networks made by combining the reactions from ℛ satisfy this principle. The number of possible reaction networks formed by choosing a subset of ℛ, including disconnected ones, is roughly given by 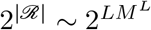. This number grows astonishingly fast with *M* and *L* (256 for *M* = 2 and *L* = 2, but ≈ 10^19^ for *M* = 2 and *L* = 4). Thus, we focus only on the *minimum* networks which can produce a given set of *N*_tgt_ target chemicals from a given set of *N*_in_ input chemicals (in metabolic terms, those that can produce biomass from the available nutrients) using the smallest possible number of reactions.

We define a *globally* minimum network to be the set of reactions ℛ _0_ ⊂ ℛ with the smallest |ℛ _0_| which can synthesize all the targets from the input chemicals. In contrast a *locally minimum* network is one from which no reaction can be removed while still maintaining production of all targets from the input chemicals. There are many locally minimum networks where the number of reactions |ℛ _0_| is not the global minimum. Note also that we must choose the input chemicals and targets such that the network consisting of all reactions in ℛ satisfies the condition of being able to synthesize the targets from the input chemicals, otherwise no subset will be able to do so. However, this ‘fully connected network’ is typically very far from being locally minimum, let alone globally minimum. Finding globally minimum reaction networks, where |ℛ _0_| is the global minimum, is computationally challenging. Here we exploit the fact that this computation can be represented in the form of the 3-satisfiability (3-SAT) problem. Being an NP-complete problem, no polynomial time algorithm exists [28]. However, sophisticated 3-SAT solvers are available which work very well in practice — in particular for the present problem for the minimum networks (see Materials and Methods for more details). Henceforth, we will use the term ‘minimum network’ to mean ‘globally minimum network’, and explicitly specify it if we mean a ‘locally minimum network’.

In the following, we deal with the minimum networks generated from the fully-connected network with *M* = 2 and *L* = 6, with *N*_in_ = 2 and *N*_tgt_ = 1. The fully-connected network and examples of minimum networks are depicted in Fig.1 A and B, respectively.

### Defining the chemical reaction dynamics of the constructed networks

There are multiple choices for implementing the kinetics of a given chemical reaction network. In the present manuscript, we utilize the simplest and widely used formulation — mass-action kinetics. Consider the following reaction with *n*_*s*_ and *n*_*p*_ reactants of the forward- and backward reaction, respectively.

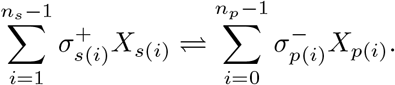

By using mass-action kinetics, the rate of the reaction (also called the ‘flux’) is given by

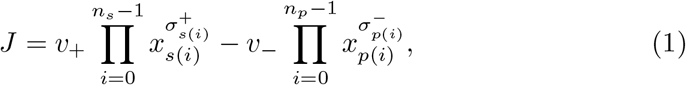

where the capital letter *X*_*i*_ indicates the *i*th chemical, and its concentration is denoted by the corresponding lower case letter. *s*(*i*) and *p*(*i*) represent the index of the *i*th reactant of the forward- and backward reactions, respectively. 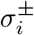 specify the stoichiometric coefficients of the *i*th chemical in the forward-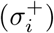 and backward 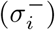 reaction, respectively. *v*_*±*_ indicates the rate of the forward (*v*_+_) and the backward (*v*_−_) reaction. When we summarize the information of the whole reaction by using a reaction stoichiometric matrix, the relationship between 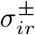(now we added the index of the reaction *r*) and the stoichiometric matrix *S* is given by

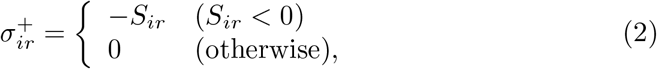

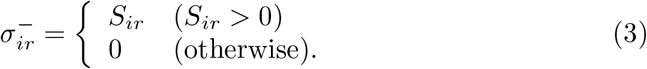

Each (minimum) reaction network is represented by a corresponding reaction stoichiometric matrix *S*_0_, which denotes the relationship of the interconversion among the chemicals by the splitting and ligation reactions. The exponents in the mass-action kinetics (Eq. (1)), 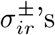, are set by *S*_0_ according to Eq. (2) and Eq. (3). This matrix however does not represent the processes of uptake of input chemicals and harvest of the targets. Therefore, we add the uptake reaction 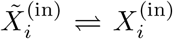 for *N*_in_ inputs, where the symbol with tilde represent the corresponding chemical in the external environment. Also, we suppose that the target chemicals are harvested together, and once harvested, the target chemicals never returns back to the reaction system, i.e, 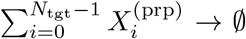. The kinetics of each of these reactions are also implemented by the mass-action kinetics given by

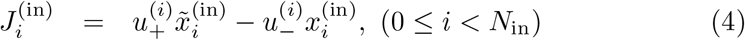

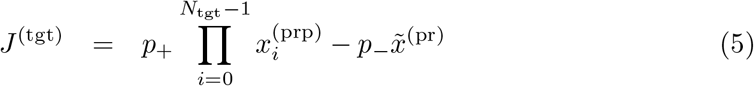

We further suppose that all chemical species are spontaneously degraded or deactivated at a uniform rate *ϕ* which is sufficiently smaller than the rates of the other chemical reactions. Additionally, the external environment is large enough so the concentrations of chemicals there are set to be constant. Then, by summing up, the dynamics of the concentrations of chemical species follow the equation below

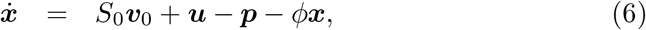

where 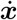 represents the time-derivative of ***x***, *d****x****/dt. S*_0_ is the stoichiometric matrix of the network. ***v***_0_ represents the reaction flux vector and each element has the form of Eq. (1). ***u*** and ***p*** denote the vectors representing the rate of input chemical uptake and target harvesting, respectively. If the *i*th chemical *X*_*i*_ is one of the inputs (target), the *i*th element of the vector ***u*** (***p***) is given by the nonzero entity ^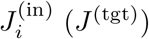^.

Once a reaction network and the inputs and targets are specified, the differential equation (Eq. (6)) is fully and uniquely determined. In this sense, we call the Eq. (6) the accompanying ordinary differential equation (ODE) of the reaction network. For simplicity, we set all the rate constants except *ϕ* to unity.

## Relaxation behaviours and network structures

In the following sections, we study the link between the network structure and the relaxation behaviour of the accompanying ODE. First, we generate ca. 2, 000 minimum reaction networks (see Methods for the parameters we used) and check if there is a steady-state attractor. As far as we have confirmed, all the networks each have a unique steady-state attractor.

For probing the relaxation characteristics, we applied perturbations to the attractor. Since the concentrations of the chemicals span a wide range at the attractors, we apply perturbations in a multiplicative manner. In addition, to normalize the strength of the perturbations, we generated the initial points on the surface of the |𝒞 _0_|-dimensional hypersphere, centred at the steady-state attractor, with a given diameter *D*. In concrete terms, a single initial point ***x***_ini_ is given by ln ***x***_ini_ = ln ***x***_att_ + *D*· ***r****/* ∥***r***∥ with ***x***_att_ is the steady-state attractor of the model, and ***r*** is a vector of the random numbers each lying in the interval [0, 1], respectively (see Materials and Methods section for more details).

For a sufficiently small value of *D*, such that the dynamics could be described adequately by linearizing the ODE around the steady-state, it is known that the relaxation dynamics would be exponential, with a rate determined by the eignevalues of the Jacobi matrix. Hence, we choose a value of *D* that is large enough to go beyond this linear regime so we can explore how the non-linearities of the ODEs result in non-exponential relaxation.

We simulated the accompanying ODE (Eq. (6)) of ca. 2, 000 minimum networks with *N*_ptb_(= 256) initial points for each network. Visually, we found three typical types of relaxation: exponential, plateau, and power-law type. Examples of each are shown in Figs.1C-F. There appeared to be three groups in terms of classification of the networks: (i) the networks exhibiting only exponential relaxation regardless of the initial concentrations, (ii) the networks showing only the plateau type relaxation regardless of the initial concentrations, and (iii) the networks displaying either the plateau or the power-law type of relaxation depending on the initial concentrations. At this stage, we do not know if the plateau exhibited by the second and the third type of networks are different in any sense, but for later arguments we refer to them as “Metastable Plateau” (independent of initial conditions) and “Confined Plateau” ^1^ (dependent on initial conditions) in Figs. 1D and E.

**Figure 1.**
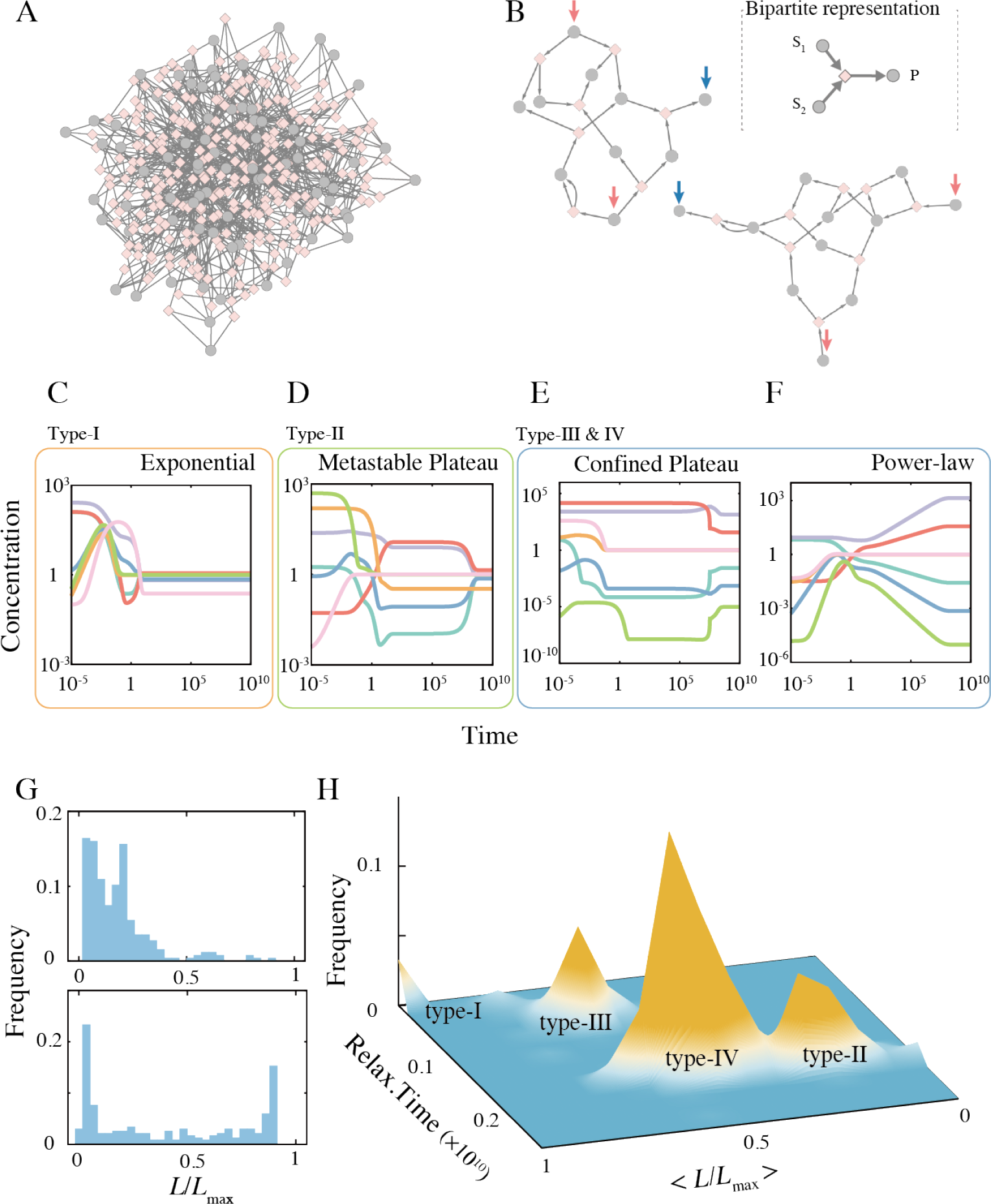
(A) The fully-connected network for *M* = 2 and *L* = 6. (B) Two examples of the minimum network which synthesizes a single target from two inputs. The inputs and target chemicals are indicated by the red arrows and the blue arrow, respectively. In A and B, the bipartite representation of the chemical reaction network is adopted. For example, the reaction *S*_1_ + *S*_2_ → *P* is depicted as follows (see inset of the panel B); first, the chemical nodes representing *S*_1_ and *S*_2_ (the gray disks) are wired to the reaction node (the pink diamond), and then, an edge connects the reaction node and the chemical node of the chemical *P* (the gray disk). Directions of the reactions are chosen to be consistent with the steady-state flux distribution. (C to F) Four typical relaxation dynamics emerged from the minimum networks: Exponential, Metastable-Plateau, Confined-Plateau, and Power-law relaxation. The vertical and horizontal axes are on a logarithmic scale. The four dynamics are classified into three category based on the networks types that exhibit the corresponding dynamics. (G) Example distributions of the migration length (see text for definition). The top panel is the distribution over all initial conditions for one network showing metastable-plateau type relaxation, and the bottom panel is from a network showing both the plateau and power-law type relaxations. (H) Classification of the minimum networks based on the average relaxation time (see text for definition) and the average migration length.

To make the classification more quantitative, we computed two quantities: the relaxation time and the migration length. Here we define the relaxation time as the time point at which the Euclidean distance, in logarithmic scale, between the state and the steady-state attractor first becomes smaller than a certain threshold value *ϵ*, that is

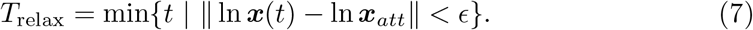

While we adopted *ϵ* = 10^−6^, it is checked that our conclusions are unchanged as long as *ϵ* value is in the range (10^−6^, 10^−2^).

Next, to distinguish between the plateau and the power-law relaxation types, we quantified the total length of the trajectory in concentration space during a given time period. It is measured by the “migration length” defined as

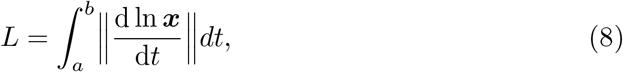

where we choose *a* = 100 and *b* = 0.01*/ϕ* in order to quantify the “motion” of the trajectory in the intermediate time scale (sufficiently larger than the time scale of the reaction rate constants, which are unity, and sufficiently smaller than that of the degradation, 1*/ϕ*). The migration length is computed in the logarithmic scale to take the effect of low-concentration chemicals into account. Additionally, we compute *L*_max_ given by Eq. (8) with *b* = 10^10^ (the end of the simulation) as the upper bound of the integral.

Our prior expectation, from visual examination of the trajectories (e.g., Figs.1C-F), was that since plateau type trajectories are almost static in the time range *t*∈ [100, 0.01*/ϕ*], the normalized migration length *L/L*_max_ would have a relatively lower value, compared to that for power-law type dynamics. We also expected that exponential relaxation dynamics will result in lower *T*_relax_ values than the other two types.

Examples of the distribution of migration length of two sets of networks are presented in Fig. 1G. The upper panel is the distribution over all initial conditions for one network that shows Metastable-plateau dynamics, and the bottom panel is for one network exhibiting both confined-plateau and power-law relaxation. The bottom distribution shows a clear double-peak due to bimodality of the relaxation dynamics, while the upper distribution is rather unimodal. We suppose that if networks exhibit a bimodal distribution due to the presence of both power-law relaxation in addition to confined-plateau relaxation, the average value of the normalized migration length *L/L*_max_ (averaged over all initial conditions) would be larger than the corresponding value for metastable-plateau networks. Thus, we utilize ⟨*L/L*_max_⟩ as an indicator of the variety of the relaxation behaviour.

The average relaxation time and the average migration length are computed for each network, and we plotted the distribution over the networks. As shown in Fig.1H, the distribution has four distinct peaks, and we classified the networks from type-I to type-IV according to the nearest peak. While the separation of the type-II and type-IV peaks is less clear, the distribution-based classification of the networks matched very well to the visual grouping of the models: the type-I networks (small *T*_relax_ and large *L/L*_max_) exhibit only exponential relaxation; most of the type-II networks showed metastable-plateau relaxation; and, the type-III networks and most type-IV networks exhibited both the confined-plateau and the power-law type relaxation depending on the initial concentrations.

Note that the final steady-state attractor is reached at the latest by *t* ∼ 1*/ϕ*, which is the timescale of the spontaneous degradation, and the qualitative differences between the different types of relaxation only appear in the *t* ≪ 1*/ϕ* region. These observations mean that the difference in the relaxation behaviour during *t* ≪ 1*/ϕ* highlight structural features of the reaction networks and would not be changed by varying the value of 1*/ϕ* as long as it is sufficiently large. What structural features of the networks result in the four different relaxation types observed in Fig.1H? We now computationally show that the relaxation behaviour is indeed determined by three features of the stoichiometric matrix (which in turn encodes the structure of the network), namely, (I) the rank gap, (II) dimension of the left null-space, and (III) the stoichiometric cone.

### I. The rank gap

The distinct relaxation behaviours are observed over timescales shorter than 1*/ϕ*, and thus, should stem from the dynamics (Eq. (6)) without the degradation term. The clear difference between the exponential relaxation and the others is the relaxation time. The exponential type trajectories already reach the attractor well before *t* exceeds 1*/ϕ*. We now show how measuring the ranks of the stoichiometric matrix and some enlargements can inform us about the reachability to the attractor in the absence of the degradation term.

First, we introduce additional notation. Let *S*_1_ denote the enlarged matrix 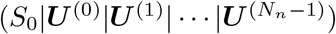, where ***U*** ^(*j*)^’s are vectors of the stoichiometry of the input uptake reactions, i.e., 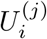 is unity if the *i*th chemical is the *j*th input chemical and zero otherwise. By vertically stacking ***v***_0_ and ***u*** in Eq. (6) and denoting it as ***v***_1_, we can rewrite Eq. (6) with *S*_1_ instead of *S*_0_. Then, removing the degradation term, we obtain

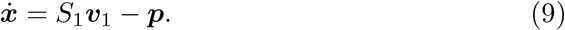

Note that Eq. (9) represents the dynamics of chemical concentration driven by the input chemical uptake, internal reactions, and the target synthesis. This equation approximates the original equation Eq. (6) for *t* ≪ 1*/ϕ*.

For there to exist a steady-state of this equation, there must be a pair of vectors, 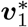 and ***p***^∗^, satisfying 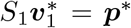. Since we are not interested in the trivial solution 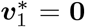, the necessary and sufficient condition for the existence of a steady-state is that the rows of *S*_1_ and ***P*** (The stoichiometric part of the harvest reaction) are linearly dependent, i.e, rank*S*_1_ = rank(*S*_1_ | ***P***), where ***P*** is the stoichiometry of the target synthesis (i.e., the *i*th element is unity if the *i*th chemical is one of the target precursors and zero for the rest).

We found that the rank gap *δ*≡ rank(*S*_1_ | ***P***) −rank*S*_1_ for all type-I networks (showing only the exponential relaxation) is zero. Interestingly, all type-II networks also satisfied the *δ* = 0 condition. On the other hand, all the type-III and type-IV networks have *δ* = 1, i.e., in the absence of degradation no steady-state exists.

This implies that there are two types of plateau relaxation. The plateaux exhibited by type-II networks are the steady-states of the model equation without degradation (Eq. (6)). However, the plateaux exhibited by type-III and IV networks are not the steady-state of Eq. (6), but are extremely slow transient dynamics towards the final steady-state for the model *with* degradation. Indeed, we find that the type-II plateaux correspond to what we had earlier termed “metastable-plateau”, while the type-III and IV plateux correspond to what we called “confined-plateau”.

### II. The left null-space

The rank gap does not distinguish between type-I and type-II networks (which both have *δ* = 0). Fortunately, these networks exhibit a unique type of relaxation regardless of the initial concentrations (exponential or metastable-plateau). Thus, we can expect that the difference stems from the network structure.

Fig.2 shows two time courses of the same type-II network but starting from different initial concentrations. The plateau regions differ not only in the concentrations of the chemicals but also in the rank order of the chemical concentrations (the identical color is used for each chemical species in both figure panels).

**Figure 2.**
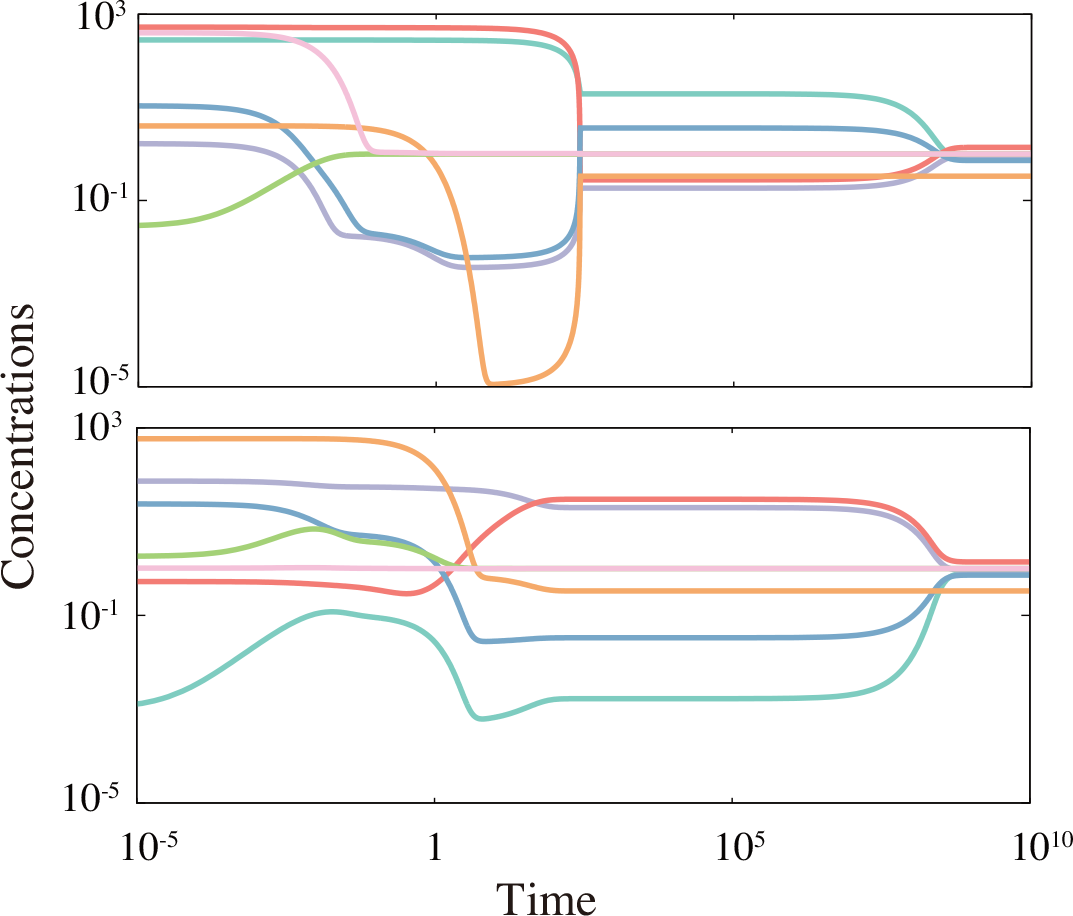
Two time courses emerged from the same minimum type-II network when started with different initial concentrations. Both the vertical and horizontal axes are plotted in the logarithmic scale.

Considering that each plateau corresponds to the steady-state of 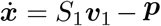 and the model eventually reaches a unique final attractor, possible mechanisms to generate the variations are limited. We hypothesized that such type-II networks might be have combinations of chemical concentrations that, over timescales much shorter than 1*/ϕ*, are constrained to be almost-conserved. If different initial conditions have different values for such conserved quantities, the system might evolve towards a different steady-state of 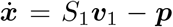 for each of these. We therefore computed the dimension of the left null-space of the enlarged stoichiometry matrix, *I* := dim (coker) *S*_2_, where *S*_2_ = (*S*_1_ | ***P***), because the left null-space of the reaction stoichiometry represents the conserved quantities of the reaction dynamics. We found that the left null-space of all the type-I models was {0}. In contrast, all the type-II models had non-zero *I*^2^. Further, we found that the type-III models had no conserved quantity (*I* = 0), while the type-IV networks had at least one (*I* ≥1).

So far, we have fully characterized the type-I and type-II networks. The type-I networks have no rank gap, *δ* = rank*S*_1_ − rank*S*_2_ = 0, and no conserved quantity *I* = dim (coker *S*_2_) = 0. Type-II networks also have no rank gap but have at least one conserved quantity, *I* ≥ 1.

### III. The stoichiometric cone

Now we move onto the networks with *δ* = 1^3^. Due to the non-zero rank gap, there is no steady-state until *t* becomes comparable with 1*/ϕ*, i.e., the spontaneous degradation starts affecting the dynamics of the concentration. Thus, in these type-III and type-IV networks, the relaxation dynamics over timescales smaller than 1*/ϕ* is not to some meta-stable almost-steady-state, but a slow transient dynamics towards the final attractor with degradation. With power-law relaxation there is a slow but continuous approach towards the final attractor, while with the confined-plateau dynamics the trajectory is practically static for a long time before suddenly “jumping” to the final attractor.

We therefore hypothesized that in the power-law case, the attractor is reachable from the initial point, but the degradation term *ϕ****x*** is necessary for making the final concentrations a stable steady-state. By contrast, in the confined-plateau case getting close to the attractor may itself be hindered in the absence of the degradation term. To check this, we introduce a subset of the phase space defined as

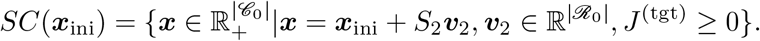

*SC*(***x***_ini_) is the region in the phase space “reachable” from the initial point by controlling the chemical reaction fluxes in any way except reversely harvesting the target chemicals. Note that in the definition, no flux balance (steady-state condition) is required. Due to the non-negativity condition (*J*^(***t****gt*)^ ≥ 0) and the non-zero rank gap *δ, SC*(***x***_ini_) does not coincide with the whole space 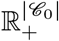. This *SC*(***x***_ini_) is equivalent to the stoichiometric cone in the field of chemical reaction network theory [29], and thus, we term *SC*(***x***) the stoichiometric cone corresponding to the initial condition ***x*** in the following arguments.

We can now mathematically formulate our hypothesis on the relationship between reachability and relaxation behaviours: for the initial points whose stoichiometric cone *SC*(***x***_ini_) contains the final attractor, relaxation becomes power-law-like, whereas, if the stoichiometric cone of the initial point does not include the attractor, plateaux appear during the relaxation (for the algorithm of judging the inclusion relationship, see Methods).

To examine the relationship between the relaxation type and the position of the initial concentrations in the phase space, we computed the average of the normalized migration length *L/L*_max_ separately for the initial concentrations whose stoichiometric cone does and does not contain the final attractor^4^. Fig. 3 is the scatter plot of the two average values for all the minimum networks of type-III and IV that we constructed. The migration length averaged over the initial concentrations with ***x***_att_ ∈ *SC*(***x***_ini_), ⟨*L/L*_max_⟩_in_, and with ***x***_att_ ∈*/ SC*(***x***_ini_), ⟨*L/L*_max_⟩_out_ are plotted in the horizontal and vertical axis, respectively. In all networks, ⟨*L/L*_max_⟩ _in_ is larger than ⟨*L/L*_max out_⟩. This result supports our intuition that some trajectories are confined in a region of the phase space and cannot get closer to the attractor.

**Figure 3.**
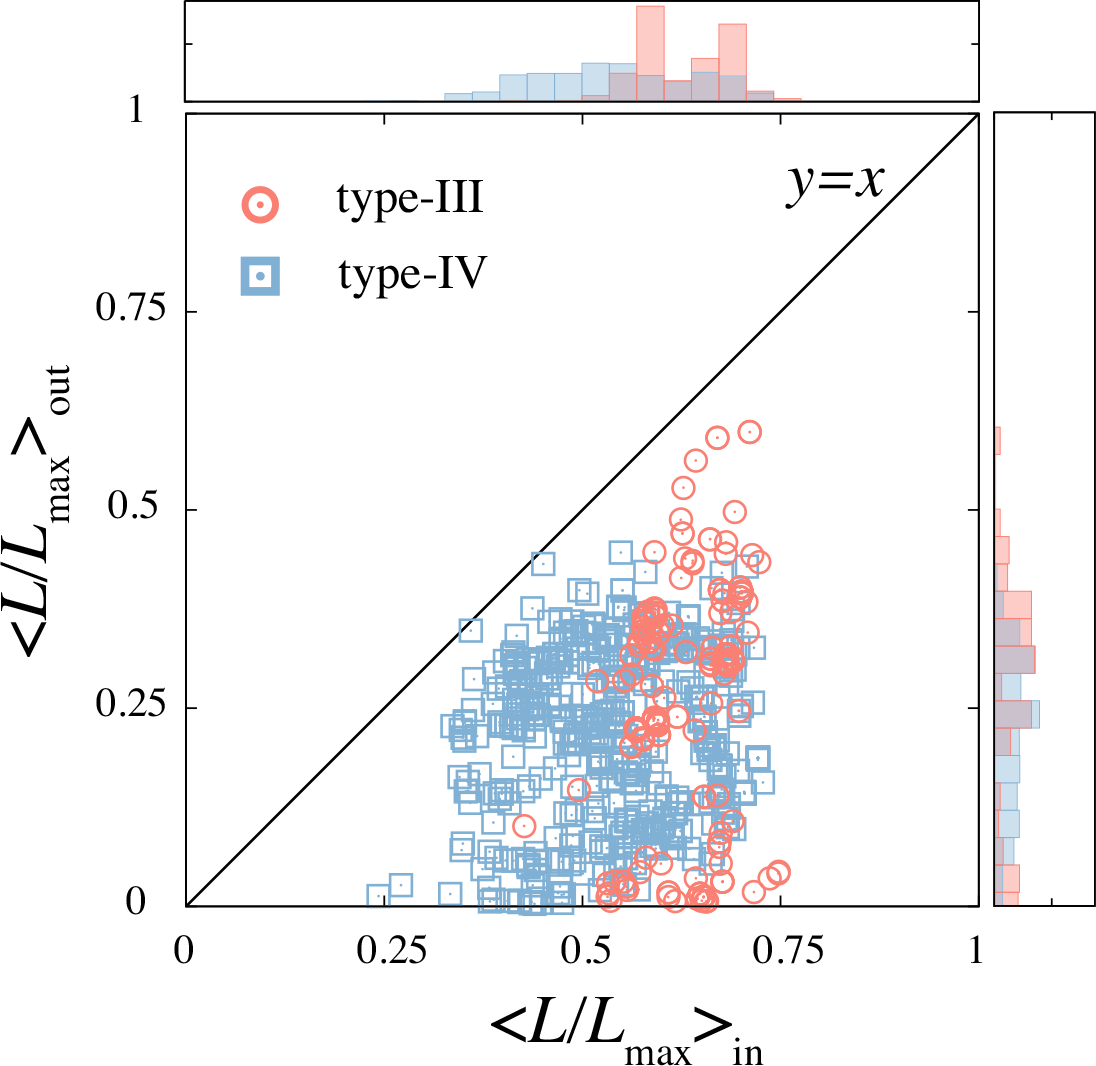
The normalized migration length is averaged separately for the time courses with the initial points whose SC, defined in the text, contains (vertical axis) and does not contain (horizontal axis) the attractor. The *y* = *x* line is plotted as a guide for the eye. The distributions of ⟨*L/L*_max_⟩ _in_ and ⟨*L/L*_max_⟩ _out_ are shown along the horizontal and the vertical axis, respectively.

The difference between the initial conditions leading to the power-law and plateau relaxation is intuitively illustrated by a simple toy model shown in Fig.4. The model equation is given by

**Figure 4.**
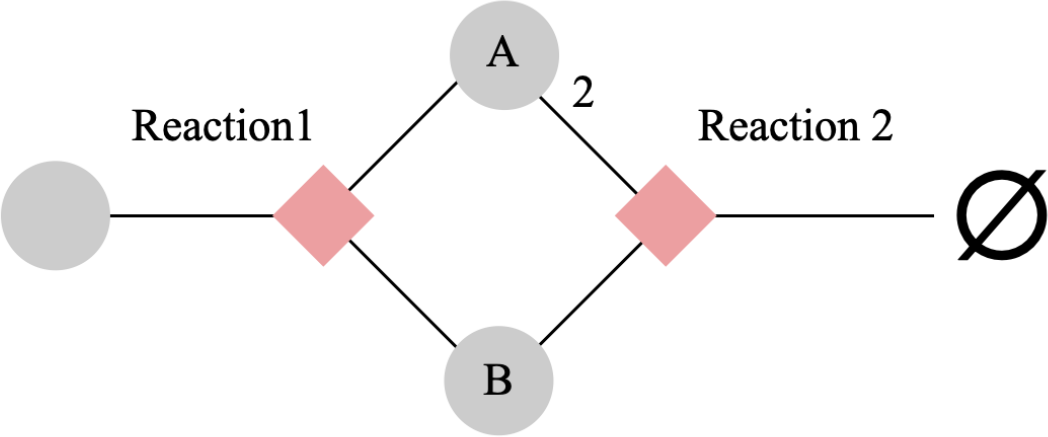
A toy model to illustrate the power-law and plateau relaxation. The leftmost chemical is the external chemical whose concentration is fixed to a constant value.

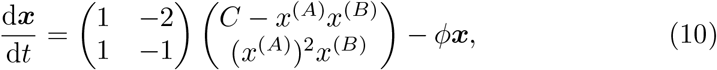

where we set the direction of the reaction from left to right on Fig.4, and *C* is the concentration of the input chemical (the leftmost chemical in the schematic). In the model, the second reaction is the target harvesting reaction.

The first reaction produces one molecule each, but the second reaction consumes two molecules of chemical A and one molecule of chemical B. As a consequence of the imbalance, the stoichiometric cone of the model is given by^5^

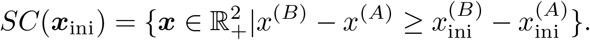

This means that the quantity *x*^(*B*)^ − *x*^(*A*)^ only increase by the two reactions. In the model, we need to consume two molecules of A to consume one molecule of B. However, to increase the concentration of A, one needs to utilize the first reaction (note that the second reaction is irreversible), increasing the concentration of B by the same amount simultaneously.

Therefore, if the initial gap of the concentration is larger than the gap at the attractor, 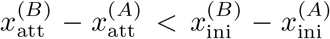, the system cannot get closer than a certain distance for *t* ≪ 1*/ϕ*. For instance, if 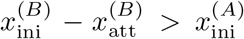 holds, the closest reacheable concentration of the chemical B is 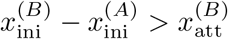^6^. In the contrast, if the stoichiometric cone contains the attractor, 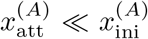and 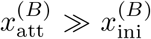 for example, the system can approach arbitrary close to the attractor even in the absence of the degradation term.

Typically, the quiescent dynamics on the confined plateau occur because the concentrations of one or a few chemicals become pretty low, as the chemical A in the toy example. Therefore, the concentrations at the plateau are not necessarily the closest possible state to the attractor in the stoichiometric cone but close to one of the axes of the Cartesian coordinate.

#### Summary of the relaxation-structure relationship

To summarize the above classification: Networks that have zero rank gap and no conserved quantity (type-I) always exhibit exponential relaxation dynamics. Networks that have zero rank gap but some conserved quantities (type-II) always exhibit plateau relaxation. Type III and type IV networks both have a rank gap of 1, but the former have no conserved quantities, while the latter have at least one. Both these networks exhibit either power-law relaxation or plateaux, depending on the initial conditions. For both types of networks, the type of relaxation for a given initial condition is determined by whether the final attractor is reachable from that initial point. If it is, then we observe power-law relaxation, and if it is not, then we observe confined plateau relaxation. Thus, from the stoichiometric matrices associated with a given network, by computing three network-structural quantities, namely, (i) the rank gap *δ*, (ii) the dimension of the left null space *I* = dim (coker) *S*_2_, and (iii) the stoichiometric cone, we can predict whether the relaxation will be exponential, power-law or plateau-type. A graphical summary of this result is presented in Fig. 5.

**Figure 5.**
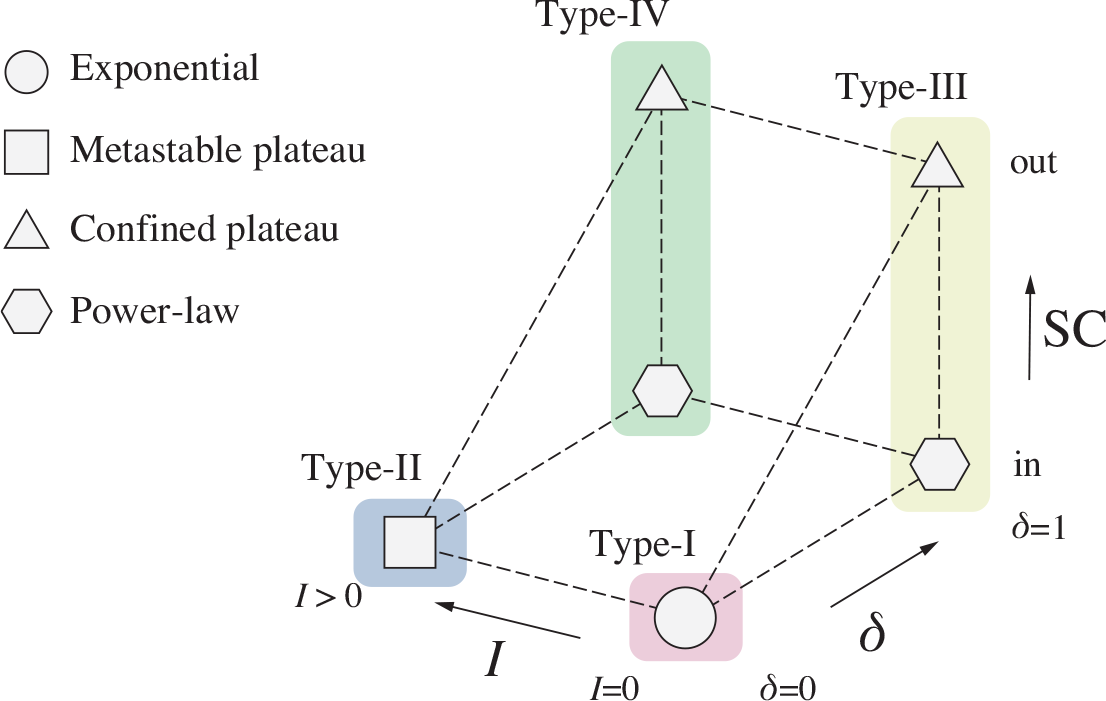
The summary of the relaxation type and network-structural features. Network types are classified according to the rank gap *δ* and the dimension of the cokernel of the stoichiometry matrix *S*_2_, *I*. In the networks with *δ* = 1, confined plateau or power-law dynamics emerges depending on whether the final attractor is in the stoichiometric cone of the initial point.

## Applicability of the structure-dynamics relationship for non-minimal networks

So far, we have explored the relationship between the network structure and emerging relaxation dynamics. There, we focused on the networks capable of synthesizing the target from given inputs with the fewest possible reactions. In this section we show that the three network-structural features predict the relaxation dynamics well in non-minimal networks also, and even in a real biological network.

### Combining minimum networks

Here we combine two minimum networks which share the same inputs and the targets. Fig. 6 is one instance of such a combination. The two networks, labelled *G* and *H*, are minimum networks that synthesize the same target from the same input. Both networks have a non-zero rank gap *δ*, and in addition, the dimension of the left-null space *I* is zero for network *G* but 1 for network *H*. The combined network is obtained as the union of the sets of chemical species and reactions of the two networks and, thus, has redundant reactions (Fig. 6A).

**Figure 6.**
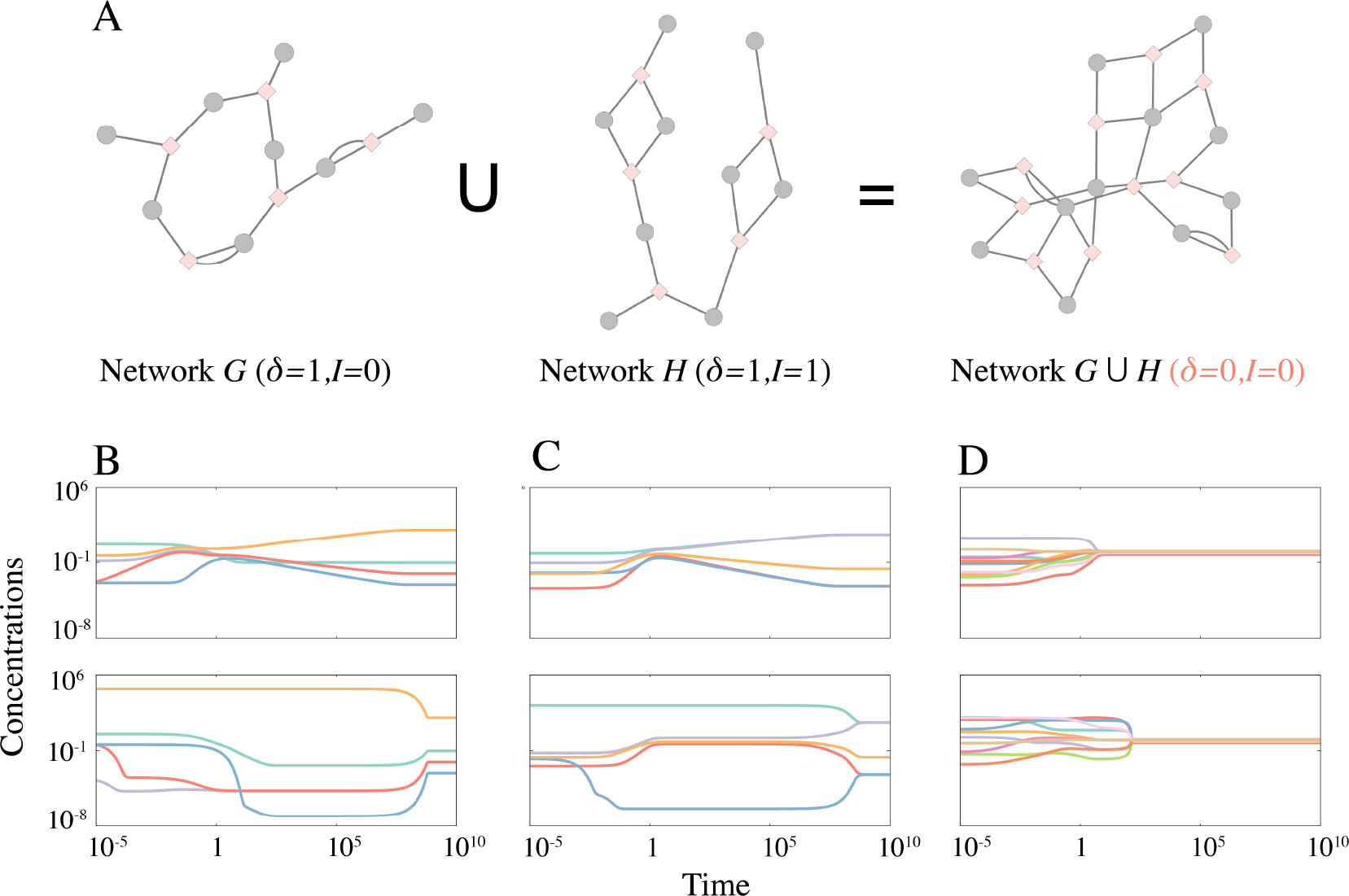
A. The two networks *G* and *H*, sharing some chemical species, including the input and the target, are combined to form the union network *G*∪ *H*. Each network *G* and *H* has a non-zero rank gap *δ*, and network *H* has one conserved quantity. However, the combined network has zero rank gap and no conserved quantity. B and C. The time courses of the network *G* (B) and *H* (C) exhibit the power-law or the confined-plateau type relaxation due to the non-zero rank gap. D. Since *δ* and *I* are both zero in the combined network *G* ∪ *H*, the relaxation time courses is of the exponential type.

Interestingly, the combined network *G* ∪ *H* has zero rank gap (*δ*=0) and no conserved quantity (*I* = 0). As a consequence, the relaxation dynamics of the combined network becomes a simple, exponential-type even though the original networks exhibit the power-law and the plateau relaxation, respectively (Fig. 6B-D).

We can also see that the effective structure of the network varies over time when we combine the two networks with different rate-constants. We construct a combined network which has rate constant unity for all reactions from network *H*, and rate constant 1*/τ* ≪ 1 for those reactions only in *G*. This combined network has an effective structure with non-zero *δ* and *I* until the dynamics start to “feel” the additional part brought from *G* (*t* ≲ *τ*). Thus, the relaxation type becomes the power-law or the plateau type for *t* ≲ *τ*. Subsequently, once the dynamics feel the whole structure of the combined network, both *δ* and *I* are zero, and thus, the relaxation dynamics transition to the exponential-type and quickly relax to the steady-state (Fig. 7).

**Figure 7.**
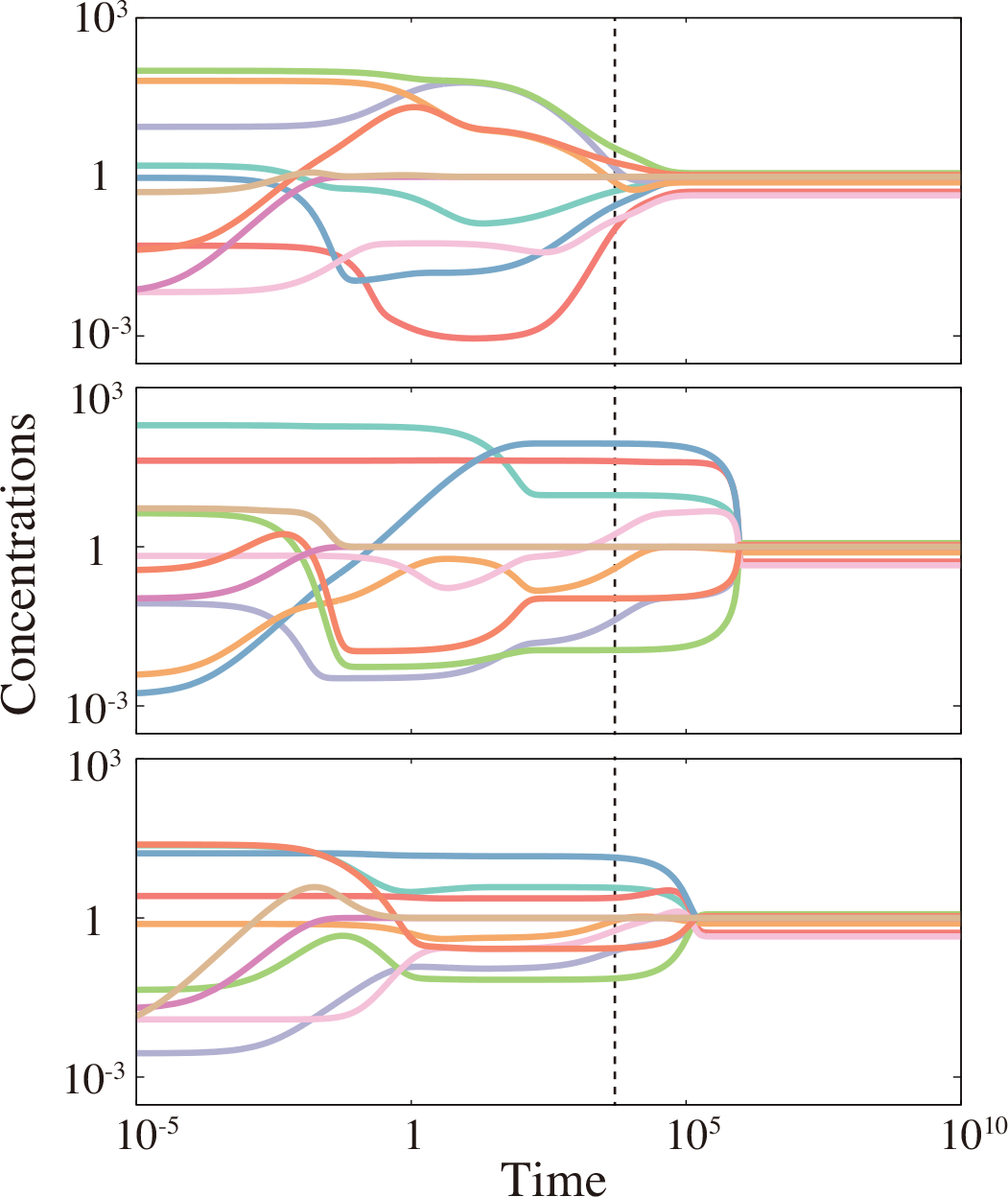
Three time courses exemplifying the dynamics of the combined network *G*∪ *H* with different rate-constants of the reactions (unity for reactions in *H* and 1*/τ* for that only in *G*). The gray-dashed vertical lines indicates *t* = *τ* = 10^4^. The dynamics for *t* ≲ *τ* shows the plateauor the power-law relaxation behaviour, while after that, the state exponentially converges to the steady-state attractor, except the left bottom case.

### Large networks

Next, we ask if the three quantities, the rank gap, the left null-spaces, and the stoichiometric cone, also work as indicators of the relaxation dynamics of large networks. For this purpose, we generated a number of reaction networks consisting of a specified number of chemicals and reactions utilizing Mixed Integer Linear Programming (for details, see Methods). In the following, we deal with |𝒞 _0_| = |ℛ _0_| = 64 networks constructed from the fully-connected network with *M* = 2 and *L* = 8 model. We chose the number of the input chemicals *N*_in_ as 2 and the number of target chemicals *N*_tgt_ as 1.

We classified the networks based on the rank gap (*δ*) and the dimension of the left null space (*I*) of the stoichiometric matrix. Examples of the dynamics of type-I and type-II networks are shown in Fig. S1. In line with the analysis of the minimum networks, most of the type-I and type-II networks exhibited the exponential-, and plateau-type relaxation, respectively, regardless of the initial concentrations. However, when the concentrations of the chemicals at the attractor span a wide range, several reactions having chemicals with low concentrations are considerably slowed down, and the relaxation time course cannot be described simply by the words “exponential” or “plateau”. The critical values of the concentration range are studied in Fig. S2.

The relaxation dynamics get much more complicated for the networks with a non-zero rank gap *δ >* 0. Three example time courses are shown in Fig. S3. Some trajectories are far from what we can call “power-law” or “plateau” relaxation. However, since the migration length is still capable of quantifying the differences of the relaxation dynamics, we examine the relationship between the migration length and the confinement of the trajectory in the stoichiometric cone.

Fig.8 is the scatter plot showing the relationship between ⟨*L/L*_max_⟩_in_ and ⟨*L/L*_max_⟩_out_ of each large network we constructed. In contrast to the minimum network case, the relationship ⟨*L/L*_max_⟩ _in_ *>* ⟨*L/L*_max_⟩ _out_ is violated by a few large networks, but the broad trend remains the same – the trajectories confined in the stoichiometric cone tend to show lower values of normalized migration length.

**Figure 8.**
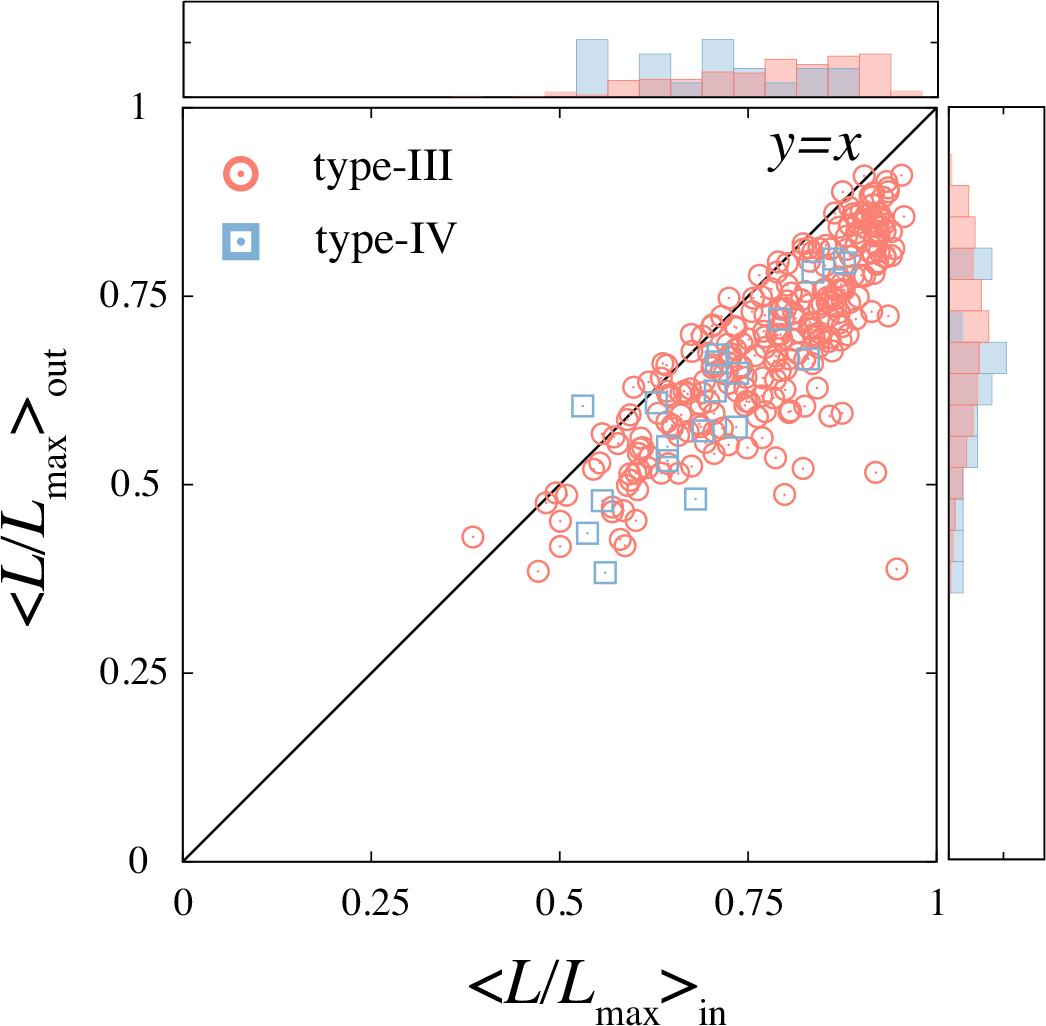
The normalized migration length is averaged separately for the time courses with the initial points whose SC contains (vertical axis) and does not contain (horizontal axis) the final attractor. The *y* = *x* line is plotted as a guide for the eye. The distributions of ⟨*L/L*_max_⟩ _in_ and ⟨*L/L*_max_⟩ _out_ are shown along the horizontal and the vertical axis, respectively.

### A biological example

Lastly, we demonstrate that the features of the network structure that we have highlighted, also play a role in a real biological network, namely, the metabolic network model of glucose fermentation in *Lactococcus lactis* (Fig.9A). The model is adopted from the reference [30]. The network takes up glucose (GLC) from the environment and ferments it. The target of the fermentation — either lactate (LAC), ethanol (ETOH), acetate (AC), acetoin (ACET), or butanediol (BUT) — is released into the environment. We briefly note some details of the model: the reaction rates are given by the Michaelis-Menten type functions [31], and the spontaneous degradation term (− *ϕ****x***) is not implemented in the original model. The reactions depicted in the network diagram are all the reactions in the model (for a full description of the model, see the original paper).

**Figure 9.**
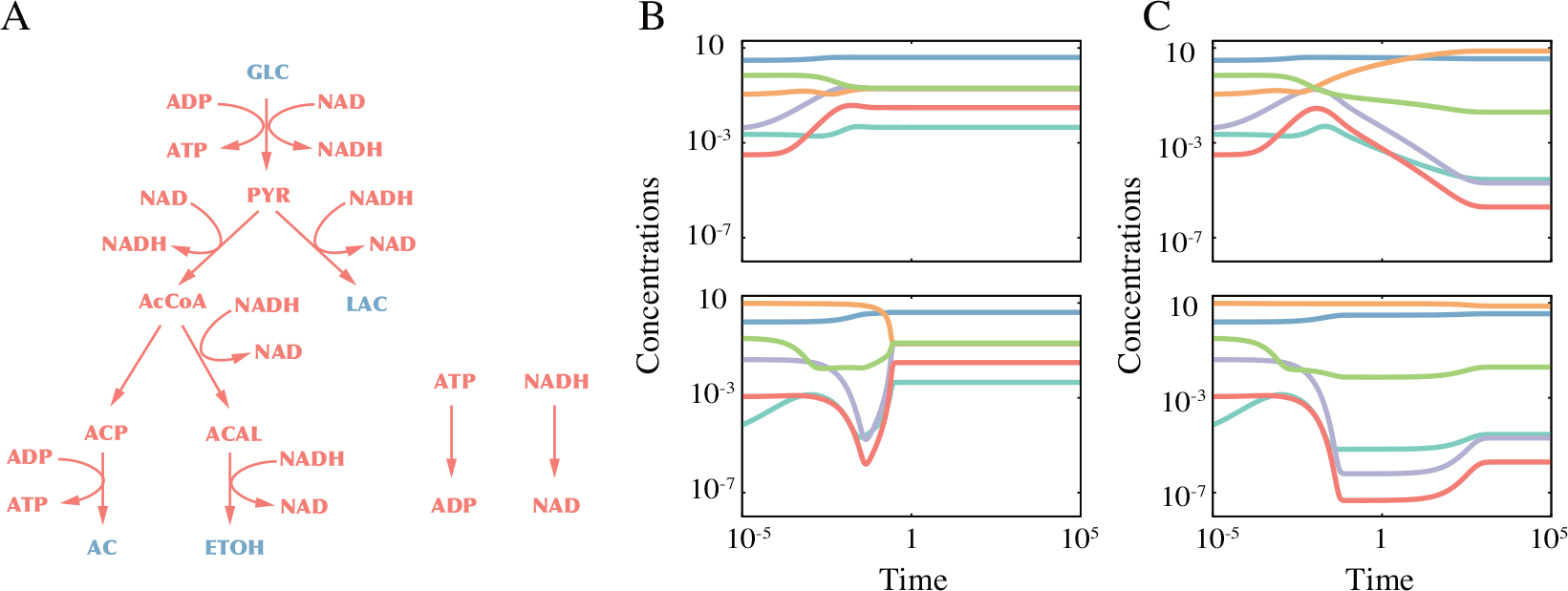
(A) The metabolic model of glucose fermentation in *Lactococcus lactis* adopted from [30]. The red colored metabolites and the blue colored metabolites are the intracellular- and extracellular metabolites, respectively. (B and C) The representative time courses of the model with the fast NOX reaction (B) and the slow NOX reaction (C). The time courses shown in the neighboring panels start from the same initial concentrations.

A significant feature of the model is the balance between NAD and NADH. Suppose that a single glucose molecule is converted into either lactate or ethanol, then the production and consumption of NAD and NADH each balance. However, when a glucose molecule is converted to acetate and secreted, a single NAD molecule is consumed, and a single molecule of NADH is produced. Thus, if the pathway GLC → AC is utilized, the concentrations of NAD and NADH cannot balance without the NADH oxidase (NOX) reaction (NADH → NAD) drawn separately from the main network in Fig.9A. Indeed, the stoichiometric matrix of the fermentation model will have a rank gap *δ* = 1 when NOX is removed from the model, while with this reaction included it has a zero rank gap.

Here, we expect to observe a qualitative difference in the relaxation dynamics that originate from the rank gap by comparing the relaxation dynamics of the model with- and without the NOX reaction. However, the model without the NOX reaction can never reach a steady-state attractor. Recall that the relaxation to the attractor in the polymer models is possible because of the spontaneous degradation. We cannot introduce it into this model because the total concentrations of the following chemical pairs are set to be constant: ATP and ADP, NAD and NADH, and Coenzyme-A (CoA, which does not appear in the network diagram) and Acetyl-CoA (AcCoA). Instead, we changed the rate constant of the NOX reaction, 100 times faster and slower than the value used in [30], which results in the effective structure of the network in a certain time range to be with- and without the NOX reaction. Here, the slow down of the NOX reaction corresponds to the decrease of oxygen in the medium. Also, we set the external acetate concentration to be 10^−4^ (mM), leaving the rest of the parameter values unchanged.

Figs. 9B and C show the examples of the relaxation time course with the fast (B) and slow (C) rate constant of the NOX reaction. As seen from the figures, the relaxations are exponential-like in the model with a fast NOX reaction, and are power-law or plateau-like in the model with the slow NOX reaction.

## Discussion

In the present manuscript, we have studied a class of chemical reaction networks consisting of polymer chemicals that can split and ligate with each other. The networks satisfy the law of mass conservation and can generate a variety of mass-conserving sub-networks. The (globally) minimum networks capable of synthesizing a specified set of target precursors from a set of input chemicals are computed within a reasonable time by utilizing the algorithm for solving the SAT problem.

We generated a large number of such minimum networks and simulated their chemical reaction dynamics to unveil the relationship between the network structure and the relaxation dynamics after perturbation of the steady-state. We found three types of relaxation dynamics: exponential, power-law and plateau type. In the first instance it is interesting that even though we choose all reaction rates to be unity, i.e. we do not introduce by hand multiple timescales into the system, much longer timescales emerge spontaneously in the relaxation dynamics. In this simplified setting therefore such slow dynamics or long-term memory must arise from the network structure. We found that we could predict which relaxation behaviour will be exhibited from three structural features of the reaction networks, namely, (i) the rank gap, (ii) the dimension of the left null space, and (iii) the stoichiometric cone. We showed that these three features determine the relaxation behaviour not only in the minimum networks, but also in combinations of the minimum networks, in large networks with many redundant pathways, as well as in one example of a real metabolic network.

Our results therefore provide easily computable criteria for predicting whether a given chemical reaction network can lead to slow dynamics. By examining the rank gap and the left null-space, one can predict whether the concentration of the chemicals will quickly relax to a steady-state or not. A non-zero rank gap indicates that the reaction network cannot have a steady state in the intermediate timescale. However, as illustrated by the network combination example and the metabolic networks of *Lactococcus lactis*, the behaviours of the chemical reaction network reflect the effective network structure at the timescales of interest. Therefore, over certain timescales, metabolic, and other real chemical networks, may effectively behave as if they have a non-zero rank gap, leading to non-exponential relaxation behaviours.

Finally, we would like to emphasise the flexibility of our model framework. Even with only polymer chemical species, the network size is easily controlled by manipulating the maximum length *L* and types of the monomers *M*. Further, the types of species and chemical reactions used to construct the networks can easily be expanded without affecting key chemical properties like mass conservation. For instance, one could include all species and reactions known to occur in microbes [32]. The SAT formulation can be equally easily used in such a case also to construct minimum networks. We expect that our approach for constructing and analyzing such artificial reaction networks will provide a useful toolbox for exploring the extensive world of out-of-steady-state dynamics of chemical reaction networks.

## Method

### Generating artificial reaction networks

For the set of chemicals 𝒞, we wish to identify the smallest set of reactions ℛ _0_ ⊆ ℛ such that target chemicals can be produced from the available inputs.

As input we have the set of available input chemicals *U* ⊂ 𝒞 and the set of target chemicals *V* ⊂ 𝒞 that must be produced. The main idea of our approach is to consider the Boolean expression

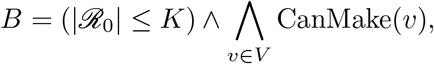

where CanMake(*v*) depends on the input chemical set *U*. We may then iteratively solve this SAT problem starting from *K* = 1 (to |ℛ |) and stop when we can satisfy the expression. This ensures that we find the global minimum solutions.

The Boolean *at most* expression, |ℛ | _0_ *K*, can be expressed efficiently using auxiliary variables [33]. What remains is the specification of the Boolean function CanMake. We can define this directly by the binary tree of reactions that leads from the inputs to the targets. For instance if the required target is *AA* and inputs are *AB* and *AC*, we could write

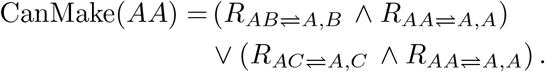

This fully defines a SAT-problem in the reaction *R*-variables. However, this approach makes exponentially long expressions for large sets of chemicals and reactions.

The reason for the long expressions is mainly repetitions: even in the tiny example above, the reaction variable *R*_*AA*⇌*A,A*_ appeared in both sub-expressions. To remedy this, we use the concept of equisatisfiability to introduce auxiliary variables that describe if a specific chemical *c* ∈ 𝒞 can be produced. Using these variables we can truncate the binary tree after one level. We thus intend to write the condition to make targets as

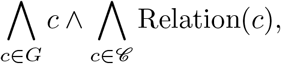

where the Boolean Relation function specifies the relationship between chemicals as specified by the set of reactions. For instance, for the above example:

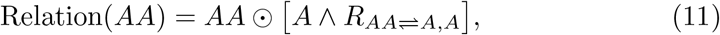

i.e. to make *AA* you need available *A* and the reaction *AA* ⇌ *A, A*. Here, ⊙ denotes *exclusive nor* indicating for *X* ⊙ *Y* that either both *X* and *Y* are True or both are False. Likewise, for example

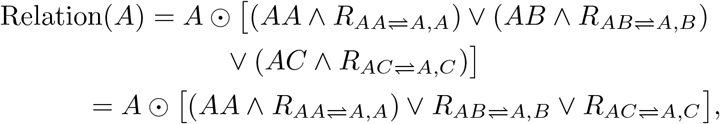

where we used the fact that *AB* and *AC* are input chemicals.

With this approach, however, we need to restrict the direction of the reactions: reactions should only be used in one direction as the network of reactions from inputs to targets should be *directed* and *acyclic* (a DAG). To fix the directionality, we introduce Boolean direction variables {*D*_*i*_}with the convention that *D* = True indicates breakdown reactions and *D* = False indicates assembly. If we denote the reaction *AA* ⇌ *A, A* by *k*, Eq. (11) thus becomes DRelation(*AA*) = *AA* ⊙ [*A* ∧ *R*_*k*_ ∧¬*D*_*k*_], whereas the DRelation for *A* would use *D*_*k*_ without the negation. When there is no reaction, we do not care about the direction of the (non-existent) reaction and we thus include a term that enforce *D*_*k*_ = True if there is no reaction, i.e. ¬ *R*_*k*_ ⇒*D*_*k*_ = *R*_*k*_ ∨*D*_*k*_.

The final reformulated expression is

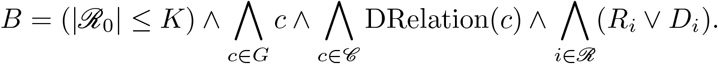

The only remaining complexity is the requirement that the solution of reactions is acyclic. We implement this requirement outside the language of Boolean expressions and simply discard such solutions. If only cyclic solutions are found at a certain *K*, these solutions are disallowed at larger *K*, thus avoiding a large number of degenerate cyclic solutions at large *K*.

All solutions to the SAT-problem are found using the Glucosamine solver via the boolexpr interface.

### Checking if the attractor is in the stoichiometric cone

For checking if the attractor is in the stoichiometric cone of the initial point, *SC*(***x***_ini_), we computed the *L*^∞^ norm between the attractor and the points in the stoichiometric cone and minimize the norm, by utilizing the following linear optimization problem.

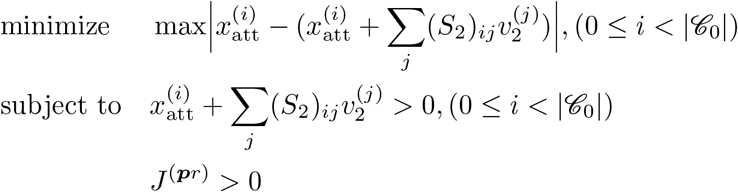

If the minimum distance is smaller than 10^−12^, we regard that the attractor is in the stoichiometric cone and *vice versa*.

### Finding 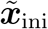

For obtaining 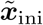, the nearest point to the initial point with the same value of the conserved quantities ***Q*** with that of the attractor, we utilized the following quadratic optimization problem with ***δx*** as variables

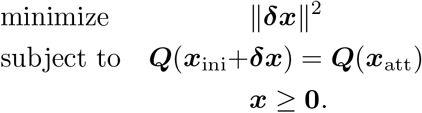

and obtained 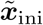 as 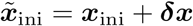.

### Generating networks with a specified size

Due to the computational cost, we adopt a mixed-integer linear (MIL) optimization approach for generating the large (non-minimal) networks, instead of the SAT approach. To obtain a network with |𝒞 _0_| chemicals, |ℛ _0_| reactions, *N*_in_ inputs, and *N*_tgt_ target precursors by solving the following MIL problem:

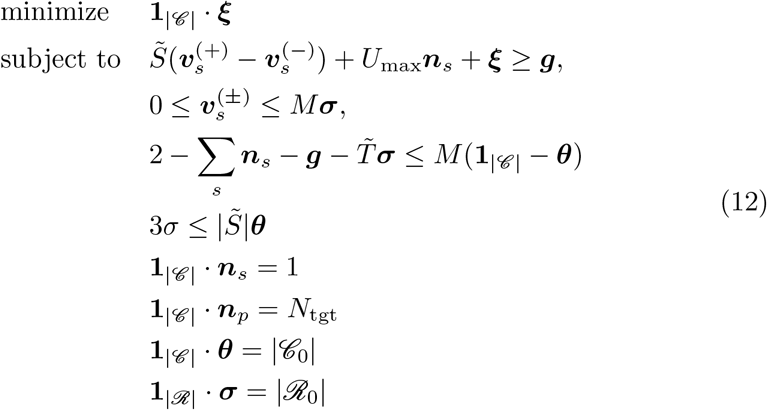

where the continuous variables are ***ξ*** ≥ 0 and 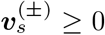, and the bool variables are ***n***, ***n***, ***θ, σ. ξ*** and 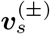 are the slack variable and the forward- and backward reaction flux vectors under the environment *s*, respectively. ***n***_*s*_ and ***n***_*p*_ denotes the choice of the input and the target. If the *i*th element of the vector ***n***_*s*_ (***n***_*p*_) is one, the *i*th chemical is the input supplied under the environment *s* (the target). ***θ*** and ***σ*** denotes which chemicals (***θ***) and reactions (***σ***) are used. If the entity is unity, it is used.

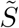 and *U*_max_ are the stoichiometric matrix of all reactions in ℛ, and the maximum uptake rate of the input chemicals. 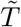 and 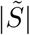 are defined as 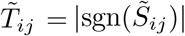 and 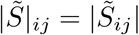. **1**_*n*_ is the abbreviated representation of *n*-dimensional vector with all the elements being unity.

Each constraint of Eq. (12) represents the following conditions. (First line) The total production rate of the *i*th chemical should be larger than *p*_*i*_ (= 1 if it is one of the targets and = 0 otherwise). (Second line) The *r*th reaction flux can be turned-on if *σ*_*r*_ is one. (Third line) If *θ*_*i*_ can be unity only if two or more (one or more for the inputs and the targets) reactions connected to the *i*th chemical are turned-on. (Forth line) To turn the *r* th chemical on, the inputs and the target of the reaction should be in the network. The fifth to the eighth line set the number of the inputs, targets, total chemicals, and total reactions.

Eq. (12) is always feasible and if the optimal value of the objective is smaller than 10^−12^, the original network {𝒞, ℛ} has a subnetwork of the specified size. For generating multiple networks, we introduce an additional constraint

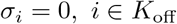

where the index set *K*_off_ consists of randomly-chosen indices.

## Acknowledgments

The authors thank Chikara Furusawa for fruitful discussion. This work is supported by the research grant (00028054) from VILLUM FONDEN and JSPS KAKENHI Grant Number JP 21K20626 and JP 22K15069. J.B.K. acknowledges funding from the Novo Nordisk Foundation, Grant Agreement NNF20OC0062047. N.M. acknowledges funding from Novo Nordisk foundation, Grant Agreement NNF21OC0068775. S.K. acknowledges funding from the Department of Atomic Energy, Government of India (Project Identification No. RTI 4006) and the Simons Foundation (Grant No. 287975).

## Footnotes

1 ”Confined” here means two things: the first meaning is that the initial conditions leading to the plateau behaviour are confined in a certain region in the phase space. The second meaning is the confinement of the trajectory in the so-called “stoichiometric cone”. See the subsection III for more details.

2 In the present minimum reaction models, there is a lower limit of *I* for the models with rank*S*_1_ = rank*S*_2_, given by *M* −*N*_*n*_, where *M* is the number of monomer types and *N*_*n*_ is the number of input chemicals. For more detail, see SI Text

3 Since the target formation reaction is represented as the single reaction (Eq. (5)), the rank gap between *S*_1_ and *S*_2_ is one at most.

4 Since the type-IV networks have conserved quantities, the stoichiometric cone of an initial point never contains the attractor unless the values of the conserved quantities at the initial point and the attractor are equal. This situation never happens by applying random perturbations to generate the initial points. Thus, we first search for the nearest point from the initial point,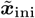, that has the same values as the conserved quantities with the attractor (see Materials and Methods). Then, we checked if the attractor is in the stoichiometric cone of 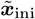.

5 Note that the stoichiometic matrix *S*_2_ for computing the stoichiometic cone in this model is 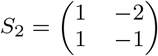

6 By inversely running the first reaction.

